# Generation of pulmonary neuro-endocrine cells and tumors resembling small cell lung cancers from human embryonic stem cells

**DOI:** 10.1101/261461

**Authors:** Huanhuan Joyce Chen, Asaf Poran, Arun M. Unni, Sarah Xuelian Huang, Olivier Elemento, Hans-Willem Snoeck, Harold Varmus

## Abstract

**SUMMARY:** By blocking an important signaling pathway (called NOTCH) and interfering with expression of two tumor suppressor genes in cells derived from human embryonic stem cells, the authors have developed a model for studying highly lethal small cell lung cancers.

**ABSTRACT:** Cell culture models based on directed differentiation of human embryonic stem cells (hESCs) may reveal why certain constellations of genetic changes drive carcinogenesis in specialized human cell lineages. Here we demonstrate that up to 10 percent of lung progenitor cells derived from hESCs can be induced to form pulmonary neuroendocrine cells (PNECs), the putative normal precursors to small cell lung cancers (SCLCs), by inhibition of NOTCH signaling. By using small inhibitory RNAs in these cultures to reduce levels of retinoblastoma (RB) protein, the product of a gene commonly mutated in SCLCs, we can significantly expand the number of PNECs. Similarly reducing levels of TP53 protein, the product of another tumor suppressor gene commonly mutated in SCLCs, or expressing mutant *KRAS* or *EGFR* genes, did not induce or expand PNECs, consistent with lineage-specific sensitivity to loss of *RB* function. Tumors resembling early stage SCLC grew in immunodeficient mice after subcutaneous injection of PNEC-containing cultures in which expression of both *RB* and *TP53* was blocked. Single-cell RNA profiles of PNECs are heterogeneous; when RB levels are reduced, the profiles show similarities to RNA profiles from early stage SCLC; when both RB and TP53 levels are reduced, the transcriptome is enriched with cell cycle-specific RNAs. Taken together, these findings suggest that genetic manipulation of hESC-derived pulmonary cells will enable studies of the initiation, progression, and treatment of this recalcitrant cancer.

## Introduction

Cancers presumed to arise from different cell lineages display characteristic genotypes, but cells of origin are generally uncertain and the relationships between lineage-specific attributes and genotypic differences of tumors are not understood (Cancer Genome Atlas Research et al., 2013; Garraway and Lander, 2013). One of the main obstacles to greater knowledge about these relationships is the need for tractable systems that allow molecular changes observed in mature cancer cells to be evaluated for their contribution to hallmarks of neoplasia as they occur during the development specific cell lineages.

Small cell lung cancer (SCLC)—,the most aggressive type of lung cancer, characterized by a poor prognosis, the rapid development of resistance to treatment, and nearly universal loss of function of tumor suppressor genes *TP53* and *RB*—is a relatively common human cancer in need of improved model systems (Gazdar et al., 2017; George et al., 2015; Peifer et al., 2012; Pietanza et al., 2015; Semenova et al., 2015). Studies of mouse models of SCLC indicate that the target cells for malignant transformation are likely to be pulmonary neuroendocrine cells (PNECs) (Linnoila, 2006; Song et al., 2012; Sutherland et al., 2011), consistent with the morphology of the cancer cells and expression of neuroendocrine markers. Despite these recent advances, fundamental features of SCLC, especially its initiation, progression, and eventual resistance to therapy, are not understood in relation to its observed genotypes.

To study these problems, we have sought ways to assess functional changes that occur after specific genes are altered in human pulmonary cells at defined stages of tissue development. Recent advances in the induction, cultivation, and directed differentiation of human embryonic stem cells (hESCs) provide opportunities to study carcinogenesis in many human cell types derived from a variety of lineages (Funato et al., 2014), including cancers such as SCLC, in which rapid onset and progression limit the availability of clinical samples, especially from early stage disease (Pietanza et al., 2015). In this report, we demonstrate that lung progenitor cells derived from hESCs can be induced to form PNECs by inhibition of NOTCH signaling, that the proportion of PNECs can be specifically increased by inhibition of expression of the *RB* tumor suppressor gene, and that subsequent interference with the *P53* tumor suppressor gene allows xenografted cells to form early stage tumors resembling SCLC.

## Results

### Generation of PNECs from cultured hESCs

Methods have recently been described for generating most, but not all, of the cell types observed in adult lung tissues by using growth factors and chemicals to alter signaling pathways sequentially in cells derived from hESCs over several weeks (Fig. 1a). Using a protocol developed by Huang et al (Huang et al., 2015; Huang et al., 2014), we have confirmed that, by day 3, about 90% of hESCs (the RUES2 and ES02 lines) differentiate into definitive endoderm (DE), triple positive for the markers KIT, EPCAM, and CXCR4 (Supplementary Fig. 1a,b); anterior foregut endoderm (AFE) by day 6; increasing numbers of lung progenitors (LP), SOX2+, NKX2.1+ and FOXA2+, between days 15 and 25 (Supplementary Fig. 1c, d, Supplementary Fig. 3a, b); and then a variety of airway and lung epithelial cells (basal progenitor cells, ciliated cells, goblet cells, club cells, and alveolar type 1 and type 2 cells (Treutlein et al., 2014; Warburton et al., 1998) [AT1 and AT2]) by day 55 (Supplementary Fig. 2a, c). However, this protocol and others produce few if any PNECs (<0.5%; Fig.1b, c; Supplementary Fig. 2b, c).

**Figure 1.**
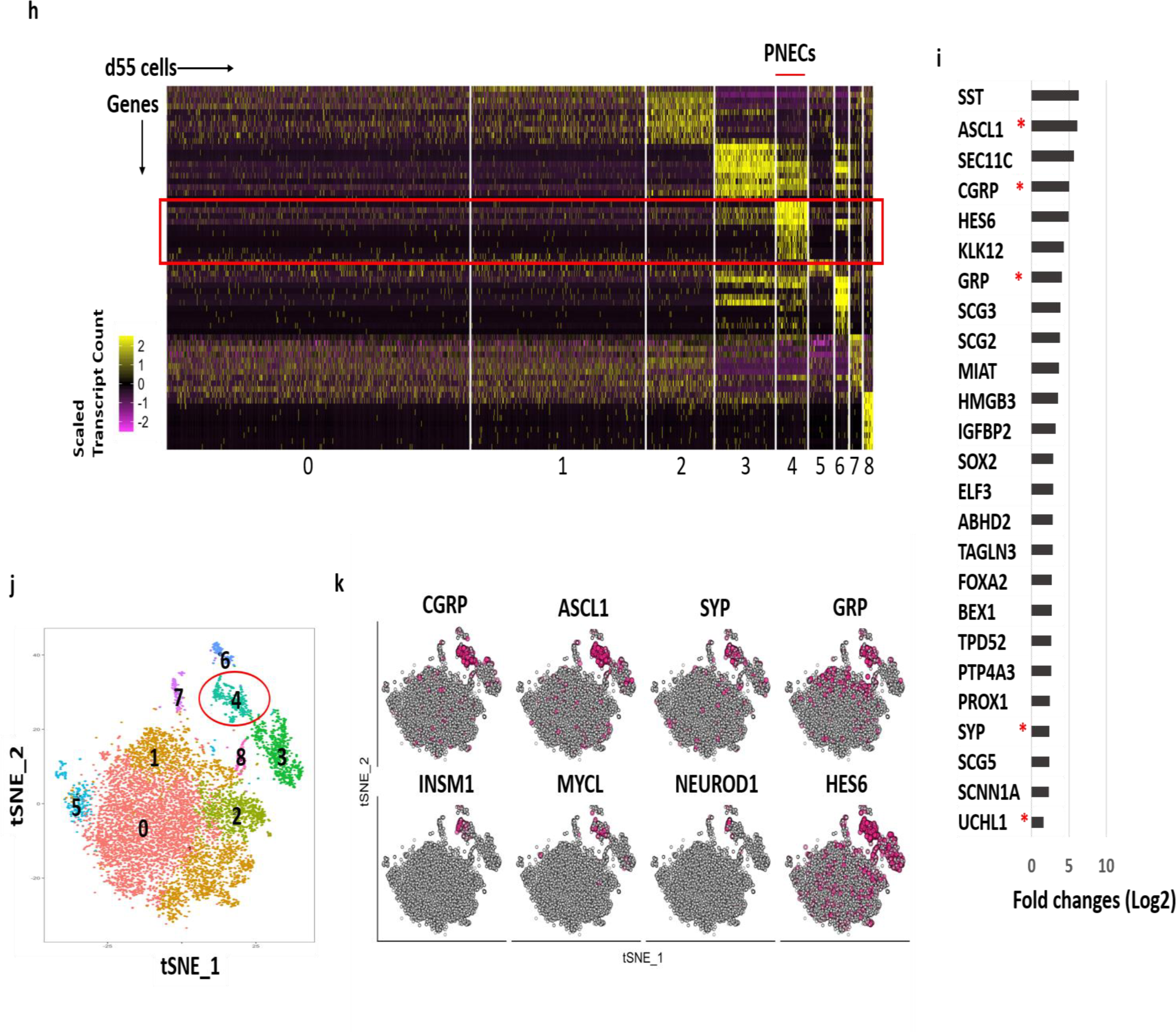
Generation and characterization of PNECs through directed differentiation of hESCs and suppression of NOTCH. **a.** Schematic of the protocol used to generate PNECs by stepwise differentiation of human embryonic stem cells (hESCs) to form definitive endoderm (DE) by day 3, anterior foregut endoderm (AFE) by day 6, and increasing numbers of lung progenitor cells (LPs) from days 15 to 25, using the differentiation mixtures I to V (defined in Methods; see Extended Data Figs 1 and 3). LPs were further differentiated in mixture VI from days 25 to 55 into the major types of lung cells (LCs) found in mature human lung parenchyma and airway epithelium (Treutlein, et.,al. 2014; Warburton, et.,al. 1998). Addition of DAPT to mixture VI induced formation of pulmonary neuroendocrine cells (PNECs; red dot), as described in the text. **b.** Detection of putative PNECs by IHC after treatment with DAPT. ESCs from the RUES2 line were differentiated according to the protocol in panel **a** to day 55 then stained to detect CGRP, NKX2.1, or both, with the indicated antisera; nuclei were detected by staining with DAPI. Scale bars: 100 µM (left) and 20 µM (right). **c.** Percentages of CGRP+ cells were determined at day 55 by FACS and displayed as flowcytometry data (red, CGRP+; yellow, CGRP-) and a scatter graph (below). **d-g.** Confirmation of mechanism of action of DAPT as inhibitor of γ-secretase cleavage of NOTCH. **d.** DAPT (5uM) treatment from day 25 to day 55 decreased level of NOTCH intracytoplasmic domain (NICD) in day 55 lung cells, as detected by western blot. **e.** LPs treated with another γ-secretase inhibitor, DBZ from day 25 to 55, also form CGRP+ cells. **f-g.** Constitutive expression of NICD prevents the appearance of CGRP+ cells co-treated with DAPT. RUES2 cells carrying a doxycycline-inducible NICD were differentiated to form LPs and then treated with DOX, DAPT, or both for 30 days. Panel **f** demonstrates the induction of NICD by DOX with Western blot; panel **g** shows by FACS that DOX (to induce expression of NICD) inhibits DAPT-mediated appearance of CGRP+ cells. ** *P* < 0.01, * *P* < 0.05 by one-way ANOVA test or (for panel **c**) by Student *t* test. Horizontal red lines denote average values; number of biological repeats (n) = 18 for panel **c;** n=10 for panel **e;** n=9 for panel **g. h-k.** Single cell RNA profiling of day 55 lung cells derived from RUES2 cells treated with DAPT (5µM) from day 25 to day 55. **h.** Heatmap representing scaled expression of the most differentially expressed genes specific to different cell clusters. Rows represent genes and columns represent cells **i.** Putative PNEC markers differentially expressed in the PNEC-like cell cluster number 4 in panel **h.** Bars indicate log fold-change versus non-PNEC cells. Asterisks indicate canonical PNEC markers. (See full gene list in Supplementary File 1.) **j.** Reduced-dimensionality t-Distributed Stochastic Neighbor Embedding (tSNE) map colored by cluster assignment (see methods). **k.** Individual cells positive for PNECs markers and other genes associated with neuroendocrine differentiation are denoted by red dots.

Studies of mouse development have suggested that inhibition of signaling via NOTCH receptors might influence cells to adopt a neuroendocrine fate (Ito et al., 2000; Linnoila, 2006; Morimoto et al., 2012; Shan et al., 2007). In addition, inactivation of *NOTCH* genes is found in about 25% of SCLCs (George et al., 2015). Based on these reports, we exposed LPs between day 25 and day 55 to N-[(3,5-Difluorophenyl)acetyl]-L-alanyl-2-phenyl]glycine-1,1-dimethylethyl ester (DAPT)(Geling et al., 2002)—a known inhibitor of γ-secretase, the protease that normally cleaves NOTCH receptors to yield a transcriptionally active, mobile, intracellular domain of NOTCH (NICD) (Schroeter et al., 1998)—to ask whether the loss of NOTCH signaling might promote the production of PNECs. (We define PNECs here as cells containing a general lung-specific marker, the transcription factor NKX2.1, and expressing one or more genes encoding well-recognized neuroendocrine markers, especially the cell surface-associated protein that includes the calcitonin gene-related peptide (CGRP) (Song et al., 2012) or the nuclear transcription factor ASCL1 (Borges et al., 1997; Borromeo et al., 2016).)

After exposure to 5 to 10uM DAPT for 30 days, a substantial number of differentiating LP cells (about 8.9 ± 1.9% in RUES2 cells, and 5.6 ± 1.4% in ES02 cells, versus 0.5 ± 0.20% or 0.4 ± 0.2% in control cultures) adopt properties of PNECs, as measured by counting CGRP+ cells with fluorescence-activated cell sorting (FACS) and confirmed qualitatively by immunofluorescence of cells in monolayer cultures with antibodies against NXK2.1 and CGRP (Fig.1b.c, Supplementary Fig. 3 c, d).

### DAPT induces PNECs by blocking cleavage of NOTCH

We took several approaches to confirm the mechanism by which DAPT induced PNECs. First, we measured the abundance of NICD in extracts from cells treated with DAPT and observed the expected loss of the γ-secretase cleavage product (Fig. 1d). To confirm the reduction in NOTCH-mediated signaling, we measured the readout from two NOTCH target genes (Iso et al., 2003), *HES1* and *HEY1,* and found a marked decrease in levels of transcripts from *HEY1* and a slight but significant decrease of RNA from *HES1* (Supplementary Fig. 4a). We also tested another known inhibitor of γ-secretase, dibenzazepine (DBZ)(Milano et al., 2004), and produced percentages of PNECs at day 55 similar to those observed with DAPT (Fig. 1e). Finally, we reversed the effects of DAPT by providing NICD, the normal product of γ-secretase-mediated cleavage of NOTCH: expression of a tetracycline-inducible transgene encoding NICD from days 25 to 55 in differentiated RUES2 cells decreased the appearance of CGRP+ cells in cultures concurrently exposed to DAPT (Fig. 1 f, g)

### Single cell transcriptional profiling of induced PNECs

To further characterize the presumptive PNECs generated by inhibition of NOTCH signaling, we used high-throughput single-cell RNA sequencing (scRNA-seq) applied to DAPT-treated and untreated cells at day 55 (Fig. 1h-k, Supplementary Fig. 2 b, c, Supplementary Fig. 4 b; see Methods). Clustering of the single-cell profiles revealed one cluster enriched with cells expressing genes that encode CGRP or ASCL1, thus identifying the presumed PNEC cells. In total, the presumed PNEC cells constituted 7.72% of the 9,824 high quality cells (pooled from two biological replicates). Analysis of differential gene expression in the various clusters revealed that cells in cluster 4 exhibit relatively high numbers of transcripts from a set of genes, including *CGRP, ASCL1, GRP, SYP,* and *UCHL1,* that encode canonical PNEC markers (Linnoila, 2006; Song et al., 2012) and other genes characteristically expressed in neuroendocrine cells (Fig. 1h, i). In comparison, we detected *CGRP* or *ASCL1* RNA in only 1.3% percent of cells, and rarely together, from control cultures not treated with DAPT (Supplementary Fig. 2c.). Investigation of the other clusters indicates that they express a variety of genes encoding markers specific for several types of lung cells, as also previously noted by Treutlein et al (Treutlein et al., 2014) (Supplementary Fig. 4b). Some of these marker genes are co-expressed in individual cells, and some are expressed in distinct populations; however, we have not pursued these observations further—for example, to propose differentiation pathways for lung development.

### Reduced expression of *RB* enlarges the proportion of PNECs

Since a central objective of this work is to assess the influence of known lung cancer genes on the behavior of cells in the lung lineage, we next examined the consequences of expressing or simulating known oncogenic mutations in hESC cultures undergoing differentiation, with or without inhibition of NOTCH signaling (Figs 2 and 3). To that end, we equipped the RUES2 hESC line with Dox-inducible transgenes encoding small hairpin RNAs (shRNAs) that inhibit production of RNA from either of the two tumor suppressor genes most commonly inactivated by mutations in SCLC, the *RB* or *TP53* genes (Supplementary Fig. 5a, b, and Fig. 3 a). We also introduced into parallel cultures inducible transgenes encoding oncogenes commonly encountered in lung adenocarcinomas, mutant *EGFR* or mutant *KRAS* (Cancer Genome Atlas Research, 2014) (Fig. 3 b).

**Figure 2.**
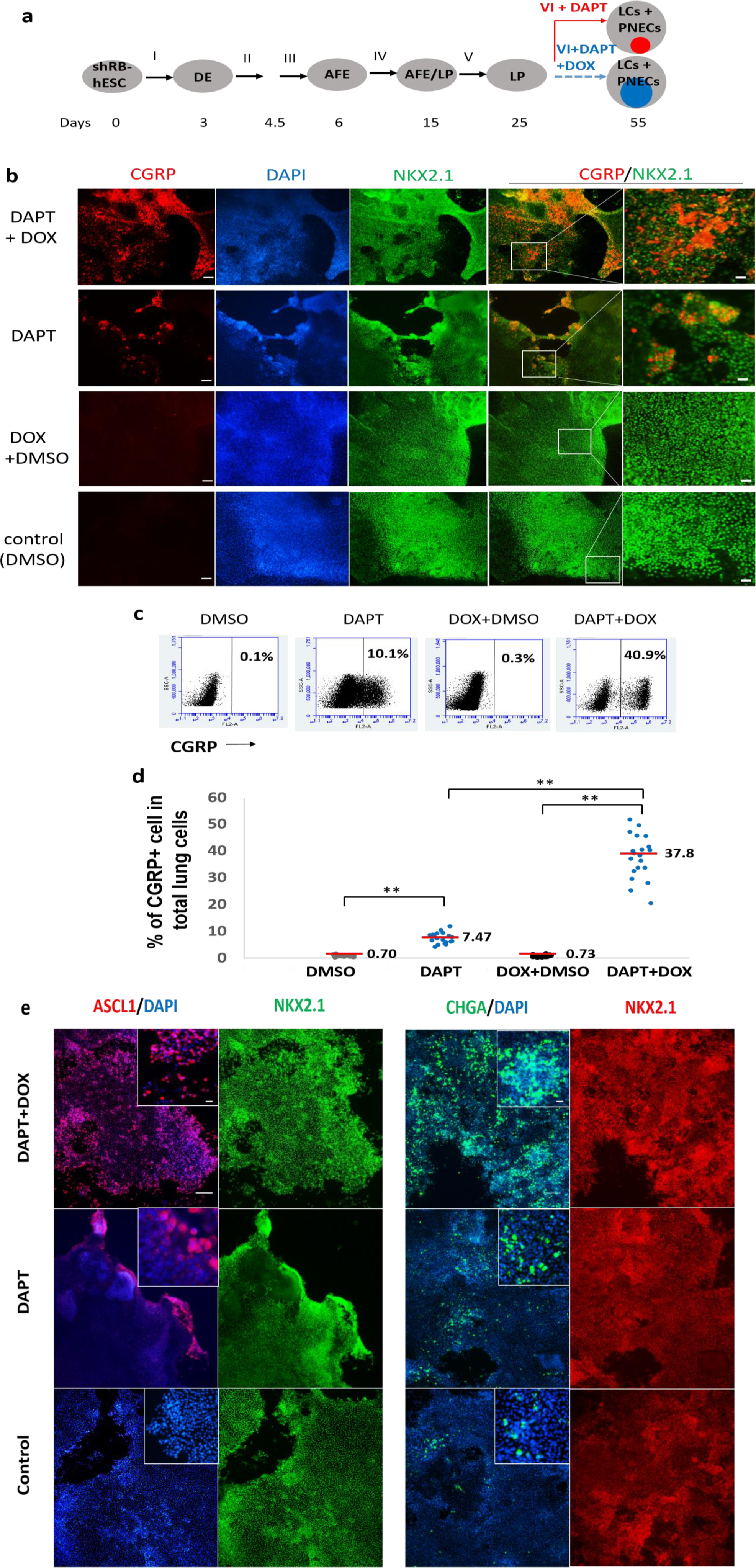
Inhibited expression of the RB tumor suppressor gene augments induction of PNECs by DAPT. **a.** Schematic of PNEC production from hESC cells carrying a tetracycline-inducible transgene that produces shRNA targeting RB1 (shRB). The format is similar to Fig.1a, except that mixture VI is supplemented with DAPT (5µM) with or without doxycycline (DOX). Internal colored circles at day 55 indicate PNECs induced by DAPT (red) or by DAPT and DOX (blue). **b-d.** Increased numbers of putative PNECs detected by co-staining for CGRP and NKX2.1 (panel **b)** or by FACS sorting with anti-human CGRP antibody (panels **c** and **d)** as described in the legend to Fig. 1 and Methods. ** P < 0.01 by one-way ANOVA test; in panel **d,** Horizontal red lines denote average values; n= 20; Scale bars: 100 µM (left) and 20 µM (right). **e.** Expression of shRNA-RB increases the percentages of ASCL1+, CHGA+ or NCAM+ cells. The indicated markers were detected by immunostaining as in panel **b.** Scale bars: 100 µM and 20 µM (in small window)

**Figure 3.**
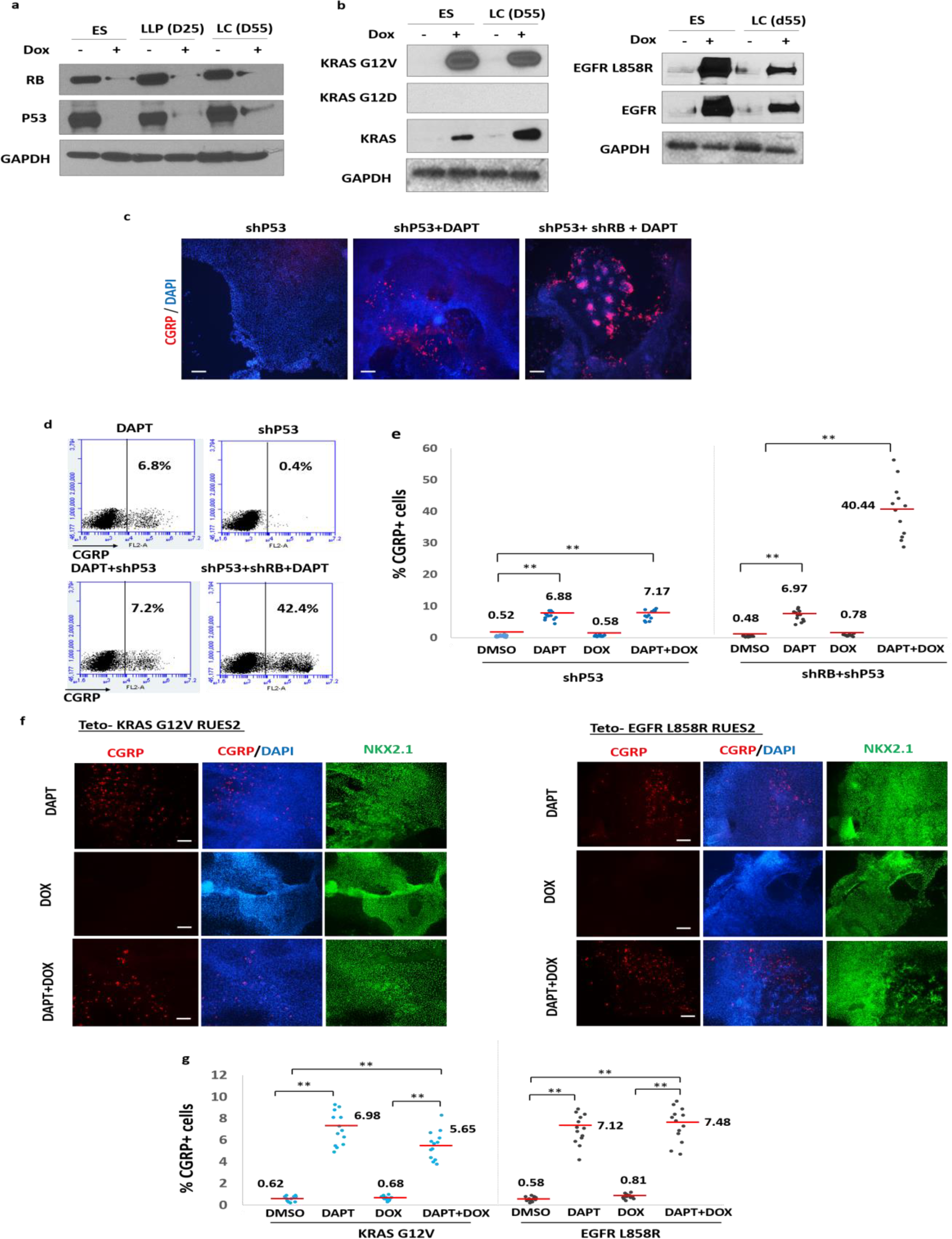
Inhibition of the p53 tumor suppressor gene or expression of two common lung cancer oncogenes do not augment production of PNECs in the presence and absence of DAPT. RUES2 cells carrying DOX-inducible transgenes that encode shRNAs targeting *P53* or *RB* or mutant alleles of *KRAS* (G12V) or *EGFR* (L858R) were used as in the experimental protocol shown in Fig.2a to measure the effects of DOX-induction of the indicated transgenes from day 25 to day 55 on production of PNECs. **a.** DOX-dependent expression of oncogenes or shRNA’s in RUES2 cells. Western blots show RB and P53 tumor suppressor proteins after DOX-induction of shRNA cassettes. **b.** Production of proteins encoded by DOX-regulated transgenes encoding KRAS-G12V (left panel) and EGFR-L858R (right panel). Proteins were detected by the indicated antibodies (see Methods). **c-g.** Decreased P53 or production of mutant KRAS or EGFR proteins fail to increase the percentage of PNECs. The indicated RUES2 cell lines were tested for the appearance of PNECs at day 55 by immunofluorescence staining for CGRP (panels **c** and **f**) and by FACS (panels **d, e, g**). See text for interpretation. Scale bars: 100 µM in panels **c;** 200 µM in panels **f,** ** P < 0.01 by one-way ANOVA test; horizontal red lines denote average values; n= 12 for **e,** and 13 for **g.**

Induction of *RB*-specific shRNA in RUES2 cells differentiating to form LCs between days 25 and 50 markedly reduced the amount of RB protein (Supplementary Fig. 5a) but not the amounts of the closely related proteins p107 and p130 (Supplementary Fig. 5b). Reduced expression of the *RB* gene was associated with a significantly increased number of CGRP+ NKX2.1+ cells (putative PNECs) from 7.5 ± 2.0% to 37.8 ± 8.2% (Fig. 2a-d), as measured by FACS, but only when cells were also exposed to DAPT to inhibit processing of NOTCH (Fig.2d). Similarly, the proportions of cells expressing the PNEC transcription factor ASCL1 and the associated markers NCAM1 and CHGA were also significantly increased (Fig. 2e).

In contrast, induction of *TP53*-specific shRNA during the same interval had no effect on the number of CGRP+ cells, with or without DAPT and with or without induction of *RB*-shRNA (Fig. 3c-e). Similarly, induction of mutant forms of EGFR and KRAS proteins between days 25 and 55 (Fig. 3b) failed to increase the number of CGRP+ cells grown in the absence or presence of DAPT (Fig. 3f, g). These findings indicate that loss of TP53 or production of mutant EGFR or KRAS proteins do not induce or affect the abundance of PNECs.

### The effects of reducing RB levels on PNEC transcriptomes

To examine the transcriptional phenotypes of cultures of differentiated (day 55) RUES2 cells in which both NOTCH and RB signaling were inhibited, we turned again to scRNA-seq. Similar clustering and differential expression analyses indicate the presence of multiple cell populations in our cultures, including an expanded PNEC-like cell compartment (11.7%), expressing markers similar to those observed in PNEC-like cells at day 55 with normal levels of RB protein (Supplementary Fig. 6e). (We attribute the relatively modest increase in PNEC-like cells as judged by scRNA-seq, compared to the increase measured by FACS, to differences in the sensitivity of detection methods that measure RNA levels as opposed to cell surface proteins.) In addition, cells in other clusters contained RNA derived from a variety of genes encoding markers specific for several types of lung cells, as also observed using scRNA-seq to examine cultures treated only with DAPT (Supplementary Fig. 7a).

When investigating the heterogeneity within the DAPT-induced PNEC cell cluster, we detected three sub-populations (Supplementary Fig.8a) with substantially different expression profiles (Supplementary Fig.8c). A similar analysis of PNECs from DAPT-treated cultures in which RB levels were reduced also revealed three subpopulations, but with transcriptional profiles different from those in cultures in which RB expression was not altered (Supplementary Fig.8a,c). Some of these differentially expressed genes, including *HES1, SST, SCG3, STMN2, ELAVL3, IGFBP4*, as well as *NEUROD1*, have previously been reported to be responsible for the heterogeneity of pulmonary neuroendocrine tumors and implicated in the initiation and progression of human SCLC(Borromeo et al., 2016; Lim et al., 2017).

To further characterize the effects of the reduction of RB protein on the transcriptional profile of the PNEC cells, we performed a differential expression analysis that compared PNECs appearing after DAPT with PNECs appearing after DAPT combined with reduced expression of *RB* (Supplementary Fig.8a). Subsequent gene function enrichment analysis (Chen et al., 2009) indicated that the most differentially expressed genes following reduction of *RB* gene expression are involved in regulation of nerve development, apoptosis, TP53 signal transduction, and other processes (Supplementary Fig.8b and Supplementary file 2).

### Transcriptomes of PNECs with reduced RB levels resemble transcriptomes from early SCLC

To ask whether inhibition of NOTCH signaling, coupled with a reduction of RB protein, produces a transcriptional program that resembles the program in human SCLC, we compared the scRNA-seq profiles of PNECs and non-PNECs from day 55 RUES2 cells, with normal or reduced levels of *RB* gene expression, to the published RNA profiles from 29 early-stage (stage Ia or Ib) human SCLCs (George et al., 2015; Peifer et al., 2012) (Supplementary Fig.7 b). This analysis confirmed that PNEC expression profiles more closely resemble SCLC profiles than do the profiles from non-PNECs. In addition, the similarity to SCLC profiles is greater (*P* < 2.2e-16 by two sided Kolmogorov-Smirnov test) in PNECs in which RB levels were reduced than in PNECs in which RB levels were not perturbed.

### Reduction of both P53 and RB in DAPT-induced PNECs allows xenografts to form tumors

To assess the ability of cells in differentiated RUES2 cultures to form tumors, we performed subcutaneous injections of day 55 cells treated in various ways into immune-deficient NSG mice (NOD.Cg-*Prkdc*^*scid*^ *Il2rg^tm1WjI^*/SzJ)(Shultz et al., 2005). The four tested cell populations all contained presumptive PNECs (measured as CGRP+ cells by FACS), ranging from about 10 to about 40 percent of the cultured cells. No growths greater than 250 mm in diameter were observed within 7 weeks after injection of parental cells exposed to DAPT alone or cells also carrying the *TP53*-shRNA or the *RB*-shRNA expression cassettes and treated with Dox (Fig 4 a). In contrast, cells with both shRNA cassettes and treated with DAPT and Dox formed visible tumors at 14 of 19 injection sites within 6 to 7 weeks. In general, these tumors were about 1 cm in diameter, formed of compact, darkly staining cells, morphologically resembling SCLC in mice and humans (Fig 4 b), and not locally invasive. The origin of the tumor cells was confirmed by detection in the RUES2 cells nuclei of GFP encoded by a component of the *RB*-shRNA cassette (Supplementary Fig. 9 b), as well as positive staining of human nuclei (Supplementary Fig. 9 c). A PNEC-like phenotype was documented using IHC to display the neuroendocrine biomarkers CGRP, NCAM1, and ASCL1, as well as the lung marker NKX2.1(Figure 4b).

**Figure 4.**
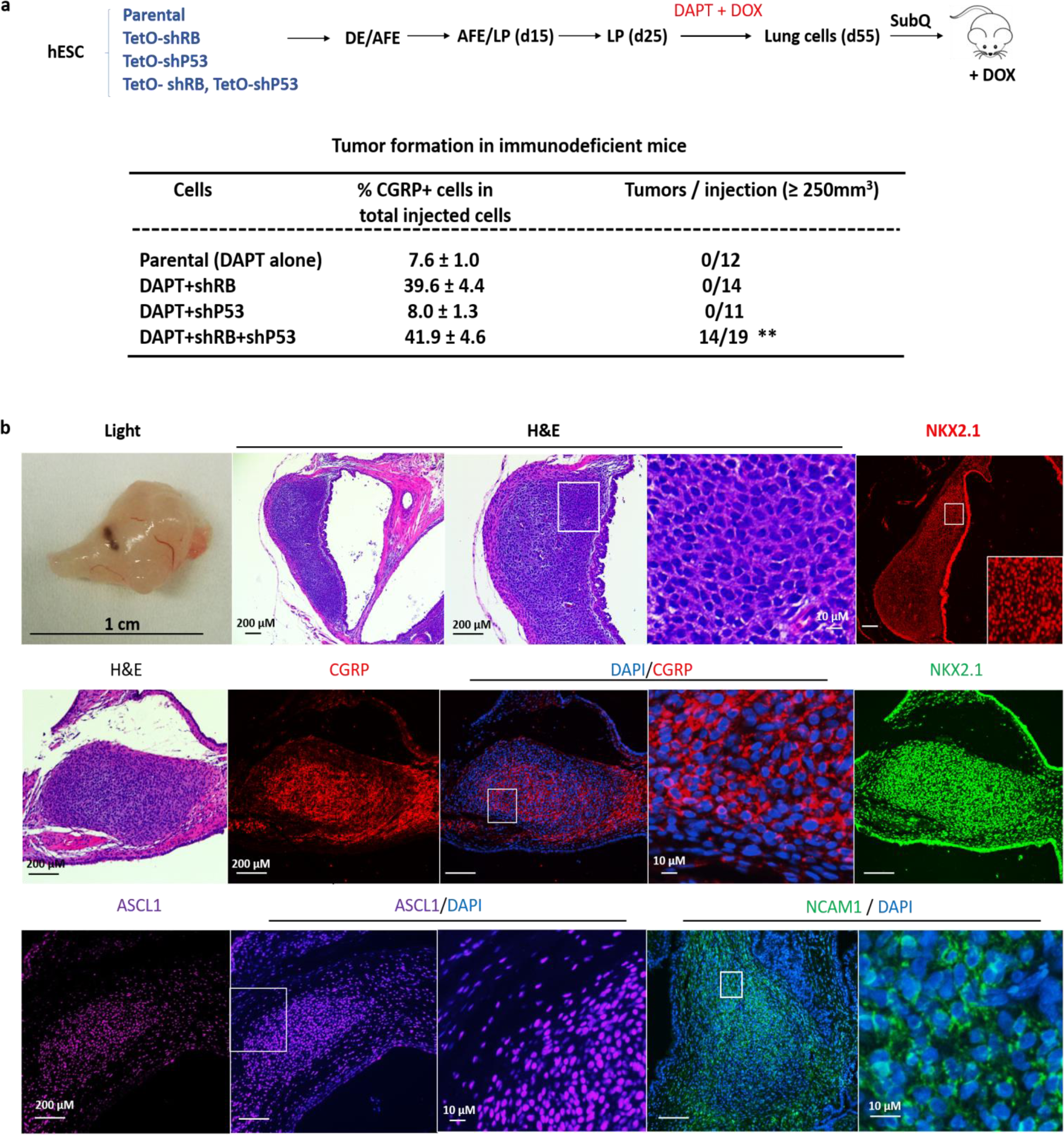
Characterization of xenografts formed with hESC - derived lung cells. **a.** Schematic representation of tumorigenesis experiment. The indicated transgenic and control lines of RUES2 hESCs were differentiated and grown in DAPT and DOX from day25 to day55. At day 55, total lung cells were injected subcutaneously into NSG mice. Xenografts grew into visible tumors after 6-7 weeks only from cells containing transgenes encoding shRNAs for both RB and P53. ** P < 0.01 by Fisher’s test. **b.** Characterization of xenografts. **Upper panel:** left segment, a representative tumor by light microscopy; middle three segments, H & E staining of that tumor at different magnifications; right segment, staining for NKX2.1; **Middle and Lower panels:** Additional samples stained with H & E, DAPI, or antibodies specific for the indicated marker proteins.

We also excluded the possibility of confusion with the teratomas known to be formed in mice injected with hESCs. Teratomas formed in NSG mice with undifferentiated RUES2 cells contained embryonic tissue markers (Liu et al., 2010), such as α-fetoprotein, Nanog, Oct4, and SSEA4 (Supplementary Fig. 9 d), and exhibited morphological features different from tumors formed with our differentiated cultures in which NOTCH, RB, and P53 pathways were disrupted (Supplementary Fig. 9a).

### The effects on PNEC transcriptomes of reduced expression of both RB and P53

Since induction of *TP53*-specific shRNA, as well as RB-specific shRNA, was required to produce tumor-forming cells in DAPT-treated cells derived from RUES2 cells, we used scRNA-seq to seek evidence that might associate the tumorigenic phenotype with changes in PNEC transcriptomes. As indicated earlier (Fig 3c-e), inhibition of expression of *TP53* does not affect the proportion of PNECs in these cultures; as expected, scRNA-seq revealed that the PNEC-like compartment of cultures in which both *RB* and *TP53* were inhibited was similar in size (10.8%) to that of cultures in which only *RB* was inhibited (Supplementary Fig. 6e). Cell clustering and analysis of differentially expressed genes in these unfractionated cultures indicate the presence of multiple cell populations (Supplementary Fig. 10 a-d), similar to cultures in which P53 was not reduced (Supplementary Fig. 6e). In the analysis of cells in which P53 levels were reduced, however, we identified two clear clusters, one differing from the other because its cells show increased expression of genes associated with active cell cycling (Supplementary Fig. 10e). This is consistent with the idea that the two clusters are actually one population of PNECs with different proliferation rates, suggesting that reduction of P53 levels, coupled with low levels of RB protein, enables a subset of PNECs to enter a proliferative mode—a phenomenon we did not observe in cells with normal levels of P53, regardless of RB status. We also found that this subset of presumptively proliferating PNECs highly expresses genes associated with inhibition of apoptosis, as well as cell cycle genes, including *BIRC5, TOP2A, MKI67, CDK1, CDKN3, CDC20*. This indicates that the main effects of reducing P53 levels in PNECs are likely to be proliferative and anti-apoptotic (Supplementary Fig. 10 e). These effects might account for the appearance of tumor-forming potential in xenografted mice.

The reduction of P53 protein in these cultures also significantly increased the expression of RNA encoding the neuroendocrine transcription factor, NEUROD1, which has been associated with advanced stages of SCLC (Borromeo et al., 2016; Osborne et al., 2013a). The percentages of *NEUROD1* RNA-positive cells in PNEC clusters produced by NOTCH inhibition alone, by NOTCH inhibition with reduced RB protein, and by NOTCH inhibition with reduction of both RB and P53, are 6.61±0.52%, 3.02±0.75% and 15.75±0.50%, respectively (mean ± sd, following repeated subsampling to account for sequencing depth). This finding is consistent with reports by several other groups that mutations in the *P53* gene promote expression of *NEUROD1* in human SCLCs (Osborne et al., 2013b).

## Discussion

In this report, we describe the development of a human cell-based model for the initiation of early stage tumors resembling SCLC. To do this, we discovered a means to induce PNEC-like cells from cultured hESCs after differentiation into lung progenitors, by inhibition of NOTCH signaling; expanded the proportion of PNEC cells by blocking the expression of the *RB* tumor suppressor gene; and conferred a tumorigenic phenotype on those cells by also blocking the expression of *TP53*, another tumor suppressor gene commonly inactivated in SCLC. We also used scRNA sequencing to characterize gene expression in hESC-derived PNECs, revealing heterogeneity in cells with normal or reduced levels of RB protein. These studies revealed the resemblance of transcriptional profiles in PNECs in which expression of *RB* was reduced to those reported for early stage human SCLC, further supporting the idea that our manipulations of hESCs in culture are generating cells on a pathway to full-fledged SCLC phenotypes. We also found that the transcriptomes of PNECs in which levels of both RB and P53 proteins were reduced include enhanced expression of genes associated with cell proliferation and inhibition of apoptosis. These changes may explain the acquisition of tumor forming potential when both tumor suppressor genes have been inhibited, as commonly occurs in human SCLC. They also indicate that it may be possible to assign specific hallmarks of the cancer phenotype to each of the two most common genotypic changes observed in SCLCs—inactivating mutations of the *RB* and *TP53* genes. Additional studies will be required to establish this idea more firmly.

Since the SCLC-like tumors grown subcutaneously in immunodeficient mice appear to have low tumor potency (slow-growing and non-invasive), it is likely that this system will enable studies of tumor progression. In addition, it should be possible to examine cells at different stages of tumor development for susceptibility and resistance to therapeutic strategies, in a manner similar to that used in an earlier study, in which a rare form of glioma was derived from hESCs by genetic manipulation of neural precursors (Funato et al., 2014).

We recognize that tumorigenesis in an intact organism is a complex process that involves interactions between the cells from which cancer arises and the several types of normal cells—stromal, epithelial, vascular, and immunological and other blood cells. These other cell types—and the adjacencies and paracrine effects that might influence tumor initiation and progression—are, of course, missing in the hESC-derived culture system we have described here. Nevertheless, this system has several advantages: it is based on the use of human rather than non-human cells, and it allows the characterization of substantial numbers of cells subjected to synchronized changes in genotype, gene expression, or hormonal effects.

Although we have begun our evaluation of the ability of manipulated lung cells to form tumors from xenografts at an ectopic site (subcutaneous tissue), it may be possible to study the cells at more appropriate locations, including lung tissue, and we are presently exploring that possibility. Other models, based on the use of genetically altered mice, offer an opportunity to study the generation of cancers like SCLC in a setting in which tissue architecture has not been perturbed, and we view the two approaches as complementary. However, we believe that the use of developmentally controlled human cells to probe the relationship between cancer genotypes and stages of lineage development to be an especially promising feature of the work we have described here.

Notably, we have generated early stage SCLC-like growths only after inducing neuroendocrine cell types and reducing expression of tumor suppressor genes implicated in human SCLC. Thus it seems likely that our differentiation protocol yields cell types, like LPs and PNECs, that are vulnerable to specific types of oncogenic lesions, such as inactivation of certain tumor suppressor genes. Further, such cells do not appear to be susceptible to the transforming effects of other genes implicated in human cancers, including oncogenes, such as mutated *EGFR* and *KRAS,* known to be drivers of other types of human lung cancer (e.g. adenocarcinomas). Understanding the determinants of this apparent specificity may help account for the widely recognized correlations between cancer genotypes and lineage phenotypes in the major forms of lung cancers and many other forms of human neoplasia (Cancer Genome Atlas Research et al., 2013; Garraway and Lander, 2013).

